# Neural representation of intraoral olfactory and gustatory signals by the mediodorsal thalamus in alert rats

**DOI:** 10.1101/2022.04.06.487193

**Authors:** Kelly E. Fredericksen, Chad L. Samuelsen

**Author notes:** **Send Correspondence to:** Chad Samuelsen, PhD, Department of Anatomical Sciences and Neurobiology, Medical Dental Research Building, Room 433, University of Louisville School of Medicine, Louisville, KY 40202.

## Abstract

The mediodorsal thalamus is a higher-order thalamic nucleus involved in a variety of cognitive behaviors, including olfactory attention, odor discrimination, and the hedonic perception of flavors. Although it forms connections with principal regions of the olfactory and gustatory networks, its role in processing olfactory and gustatory signals originating from the mouth remains unclear. Here, we recorded single-unit activity in the mediodorsal thalamus of behaving rats during the intraoral delivery of individual odors, individual tastes, and odor-taste mixtures. Our results are the first to demonstrate that neurons in the mediodorsal thalamus dynamically encode chemosensory signals originating from the mouth. This chemoselective population is broadly tuned, responds with excitation and suppression, and represents odor-taste mixtures differently than their odor and taste components. Furthermore, a subset of chemoselective neurons encoded taste palatability. Our results further demonstrate the multidimensionality of the mediodorsal thalamus and provides additional evidence of its involvement in processing chemosensory information important for consummatory behaviors.

**SIGNIFICANCE STATEMENT:** The perception of food relies upon the concurrent processing of olfactory and gustatory signals originating from the mouth. The mediodorsal thalamus is a higher-order thalamic nucleus involved in a variety of chemosensory-dependent behaviors and connects the olfactory and gustatory cortices with prefrontal cortex. However, it is unknown how neurons in the mediodorsal thalamus process intraoral chemosensory signals. Using tetrode recordings in alert rats, our results are the first to show that neurons in the mediodorsal thalamus dynamically represent olfactory and gustatory signals from the mouth. Our findings suggest that the mediodorsal thalamus is a key node between sensory and higher-order cortical areas for processing chemosensory information underlying consummatory behavior.

## Introduction

The perception of food, and ultimately the decision whether to eat it or not, requires the integration and discrimination of multisensory signals from the mouth (Sclafani, 2001; Verhagen and Engelen, 2006). While all senses contribute, the concurrent activation of the olfactory and gustatory systems is essential for giving food its flavor (Small, 2012; Prescott, 2015). This multisensory process generates enduring odor-taste associations that are crucial for guiding future food choices (Fanselow and Birk, 1982; Schul et al., 1996; Sakai and Yamamoto, 2001; Gautam and Verhagen, 2010; Green et al., 2012; McQueen et al., 2020). These experience-dependent behaviors rely upon a network of brain regions to integrate and process multimodal chemosensory signals to guide consummatory choice (Samuelsen and Vincis, 2021). The mediodorsal thalamus is a higher-order thalamic nucleus involved in an array of cognitive functions, including attention (Plailly et al., 2008; Veldhuizen and Small, 2011; Schmitt et al., 2017; Rikhye et al., 2018), valuation (Rousseaux et al., 1996; Sela et al., 2009; Tham et al., 2011; Alcaraz et al., 2018), memory (Parnaudeau et al., 2013; Bolkan et al., 2017; Scott et al., 2020), and stimulus-outcome associations (Oyoshi et al., 1996; Kawagoe et al., 2007; Courtiol and Wilson, 2016). It receives projections from primary olfactory cortical areas (e.g., piriform cortex), is reciprocally connected with the gustatory cortex, and forms dense reciprocal connections with higher-order cortical areas important for decision-making (Price and Slotnick, 1983; Kuroda et al., 1992; Ray and Price, 1992; Shi and Cassell, 1998; Kuramoto et al., 2017; Pelzer et al., 2017). As the first thalamic nucleus to receive olfactory input, studies have focused on the role of the mediodorsal thalamus in a variety of experience-dependent olfactory behaviors, including olfactory attention (Plailly et al., 2008; Small et al., 2008; Tham et al., 2009; Veldhuizen and Small, 2011), odor discrimination (Eichenbaum et al., 1980; Staubli et al., 1987; Courtiol and Wilson, 2016; Courtiol et al., 2019), and odor-reward associations (Kawagoe et al., 2007). It is implicated in the hedonic perception of odors and flavors, as people with lesions of the mediodorsal thalamus report lower hedonic ratings for experienced odors and odor-taste mixtures (Tham et al., 2011). Electrophysiological experiments in anesthetized and behaving rats show that neurons in the mediodorsal thalamus encode odors sampled by sniffing (i.e., orthonasal olfaction) (Courtiol and Wilson, 2014) and display odor selectivity during olfactory discrimination tasks (Courtiol and Wilson, 2016). However, it is unknown how neurons in the mediodorsal thalamus represent orally consumed odors, tastes, and odor-taste mixtures.

To address this question, we recorded single-unit activity in the mediodorsal thalamus of behaving rats during the intraoral delivery of three different stimulus categories: individual odors, individual tastes, and odor-taste mixtures. This approach allowed odorized stimuli to be detected via retronasal olfaction, an essential factor for the perception of flavor (Verhagen and Engelen, 2006; Prescott, 2012), and ensured that all chemosensory stimuli would share similar somatosensory and attentional attributes associated with the intraoral delivery of liquids. Our data provide novel insights into how the mediodorsal thalamus processes chemosensory signals originating from the mouth. Our findings reveal that neurons in the mediodorsal thalamus encode the sensory and hedonic properties of gustatory stimuli, demonstrate that chemosensory-evoked activity is rapid and persistent with time-varying differences between categories of stimuli, and provide evidence that odor-taste mixtures are represented differently from their unimodal odor and taste components. Together, our results suggest that the mediodorsal thalamus may be a key node between sensory and higher-order cortical areas for processing chemosensory information underlying consummatory behavior.

## Materials and Methods

### Experimental subjects

All procedures were performed in accordance with university, state, and federal regulations regarding research animals and were approved by the University of Louisville Institutional Animal Care and Use Committee. Female Long-Evans rats (~250-350g, Charles Rivers) were single-housed and maintained on a 12 h light/dark cycle with *ad libitum* food and distilled water unless specified otherwise.

### Surgery and tetrode implantation

Rats were anesthetized in an isoflurane gas anesthesia induction chamber with a 5% isoflurane/oxygen mix. Once sedated, rats were removed and placed in an isoflurane mask. Rats received preoperative injections of buprenorphine HCl (0.05 mg/kg), atropine (0.03 mg/kg), dexamethasone (0.2 mg/kg), and lactated Ringers solution (5 ml). Once a surgical level of anesthesia was reached, the scalp was shaved, and the rat was placed into the stereotaxic frame. Depth of anesthesia was maintained with 1.5-3.5% isoflurane/oxygen mix and monitored every 15 minutes with inspection of breathing rate, whisking, and toe-pinch withdraw reflex. Ophthalmic ointment was placed on the eyes and the scalp was swabbed with a povidone-iodine solution then 70% ethanol solution. A midline incision was made and the skull was cleaned with a 3% hydrogen peroxide solution. Craniotomies were drilled for the placement of 7 anchoring screws (Microfasteners, SMPPS0002). A craniotomy was made over the right mediodorsal thalamus (AP: −3.3 mm, ML: 1.4-1.6 mm from bregma) to implant a movable bundle of 8 tetrodes (Sandvik-Kanthal, PX000004) with a final impedance of ~200-300 kΩ. The medial and central portions of the mediodorsal thalamus were targeted due to the dense connectivity with olfactory and gustatory cortical areas (Price and Slotnick, 1983; Kuroda et al., 1992; Ray and Price, 1992; Shi and Cassell, 1998; Pelzer et al., 2017). The tetrode bundle was inserted at a 10° angle to avoid the superior sagittal sinus and lowered to a depth of ~4.7 mm from the brain surface. Ground wires were secured to multiple anchoring screws. Intraoral cannulas (IOCs) were bilaterally inserted to allow for the delivery of liquid stimuli directly into the oral cavity. All implants and a head-bolt (for head restraint) were cemented to the skull with dental acrylic. Injections of analgesic (buprenorphine HCl) were provided for 2-3 days post-surgery. Rats were allowed a recovery period of 7-10 days before beginning water restriction.

### Stimulus delivery and recording procedure

Following recovery from surgery, rats began a water regulation regime of 1 h access of distilled water per day in the home cage. Next, rats were given 4 consecutive days of 1 h of home cage experience with two odor-taste mixtures: a palatable mixture of 0.01% isoamyl acetate-100 mM sucrose and an unpalatable mixture of 0.01% benzaldehyde-200 mM citric acid. Rats were then trained to wait calmly in a head restrained position for the delivery of stimuli through IOCs. All stimuli were mixed with distilled water and delivered via manifolds of polyimide tubes placed in the IOCs. Stimuli included distilled water, tastes (100 mM sucrose, 100 mM NaCl, 200 mM citric acid, and 1 mM quinine), odors (0.01% isoamyl acetate, 0.01% benzaldehyde, and 0.01% methyl valerate), the previously experienced odor-taste mixtures (isoamyl acetate-sucrose and benzaldehyde-citric acid), and mismatched pairings of those mixtures (isoamyl acetate-citric acid and benzaldehyde-sucrose). These odors have been used in previous studies investigating orally consumed odors (Aimé et al., 2007; Julliard et al., 2007; Gautam and Verhagen, 2010, 2012; Tong et al., 2011; Rebello et al., 2015; Samuelsen and Fontanini, 2017; Bamji-Stocke et al., 2018; Fredericksen et al., 2019; McQueen et al., 2020). At these concentrations, isoamyl acetate and benzaldehyde lack a gustatory component (Aimé et al., 2007; Gautam and Verhagen, 2010; Samuelsen and Fontanini, 2017). A trial began with an intertrial interval of 20 ±5 s followed by the pseudo-random delivery of ~25-30 µl of water, a single taste, a single odor, or an odor-taste mixture. Each stimulus delivery was followed 5 s later by a ~40 µl distilled water rinse. All recording sessions consisted of a total of 120 trials (i.e., 12 stimuli x 10 trials), except for one session where a rat received just 9 trials per stimulus. The tetrode bundles were lowered ~160 µm after each recording session and rats were given 1 h access in the home cage to the experienced odor-taste mixtures of isoamyl acetate-sucrose and benzaldehyde-citric acid. After training days, rats were allowed 1 h access of distilled water in the home cage.

### Electrophysiological recordings

Signals were sampled at 40kHz, digitized, and band-pass filtered using the Plexon OmniPlex D system (Plexon, RRID:SCR_014803). Single units were isolated offline using a combination of template algorithms, cluster-cutting, and examination of inter-spike-interval plots using Offline Sorter (Plexon, Offline Sorter; RRID:SCR_000012). Data analysis was performed using Neuroexplorer (Nex Technologies; RRID SCR 001818) and custom written scripts in MATLAB (The MathWorks, RRID:SCR 001622).

### Analysis of single units

For each neuron, single trial activity and peristimulus time histograms (PSTHs) were aligned to stimulus presentation through the IOC. Responses were normalized using the area under the receiver-operating characteristic (auROC) (Cohen et al., 2012; Jezzini et al., 2013; Gardner and Fontanini, 2014; Samuelsen and Fontanini, 2017). This method normalizes stimulus-evoked activity to baseline on a 0–1 scale, where 0.5 represents the median of equivalence of the baseline activity. A score above 0.5 is an excited response and below 0.5 is a suppressed response. Population PSTHs are the average auROC of each neuron in the observed population. A bin size of 200 ms was used for all analyses unless otherwise specified. Neurons were defined as ‘chemoselective’ when two criteria were satisfied: (1) stimulus-evoked activity significantly differed from baseline and (2) there was a significant difference across the twelve intraoral stimuli. Significant changes from baseline were detected using a Wilcoxon rank sum test between 2 s baseline (200 ms bins) and 5 s post-stimulus delivery (200 ms bins) with correction for family-wise error (two consecutive significant bins, *P* < 0.05). Significant differences evoked by the twelve intraoral stimuli were determined using a two-way ANOVA (stimulus X time) for 200 ms bins from 0 to 5 s after stimulus delivery. A neuron significantly differed across the intraoral stimuli when the stimulus main effect or the interaction term (stimulus X time) was *P* < 0.01. Proportional analyses were performed using a *X*^2^ (*P* < 0.05). Post hoc comparisons were made using Fisher’s exact test with Dunn–Sidak correction for familywise error.

### Excited and suppressed responses

The average of the auROC normalized activity of the bins that significantly differed from baseline was used to determine whether the significant responses of chemoselective neurons were excited or suppressed. Responses whose average significant auROC score was greater than 0.5 were defined as excited, those with an average significant auROC score less than 0.5 were defined as suppressed. Comparisons in the time course between non-responses and responses excited or suppressed by chemosensory stimuli were made using the Wilcoxon rank sum test with correction for family-wise error (two consecutive significant bins, P < 0.05). The heat maps (Fig. 3B) show all the significant responses to each stimulus plotted from the lowest average significant auROC score (suppressed) to the greatest average significant auROC score (excited). Latency and duration of the significant responses of chemoselective neurons were determined using a sliding window of 100 ms, stepped in 20 ms increments until the firing rate was 2.58 standard deviations (99% confidence level) above or below the average baseline firing rate (2 s before stimulus delivery). Response latency was determined by the trailing edge of the first significant bin. Response duration was calculated as the total number of 20 ms bins significantly above (excited) or below (suppressed) the average baseline firing rate. Overall difference between excited and suppressed response latency and duration were compared using a two-sample Kolmogorov–Smirnov test (*P* < 0.05). Comparisons of response latency and duration between stimulus categories were made using the Kruskal-Wallis test corrected with the Tukey HSD test (*P* < 0.05).

### Principal component analysis

To examine the response dynamics over time, principal component analyses (PCAs) were performed in MATLAB (The MathWorks, RRID:SCR 001622) on the auROC normalized activity (−2 to 5 s; 200 ms bins) of the significant responses in each stimulus category (Narayanan and Laubach, 2009; Liu and Fontanini, 2015). Principal components (PCs) accounting for more than 5% of the variance were selected and their eigenvectors were used to describe the temporal dynamics of significant responses to each chemosensory category (Fig. 3C).

### Population decoding analysis

Population decoding analyses were performed using the neural decoding toolbox (Meyers, 2013). These analyses were used to quantify how populations of neurons in the mediodorsal thalamus represent different categories of chemosensory signals across time. For each subpopulation of neurons (e.g. chemoselective neurons vs. non-selective, mixture-selective vs. non-mixture-selective, and palatability-related vs. non-palatability), a firing rate matrix of the spike time stamps of each neuron (2 s before and 5 s after) were realigned to stimulus delivery, compiled into 250 ms bins with a 50 ms step, and normalized to Z score. Three firing rate matrices were made for each subpopulation: (1) water and the three odors, (2) water and the four tastes, and (3) water and the four odor-taste mixtures. Water was included in each category as a general non-chemosensory stimulus. A “max correlation coefficient” classifier was used to assess stimulus-related information represented by the population activity. Matrix activity was divided into 10 “splits”: 9 (training sets) were used by the classifier algorithm to “learn” the relationship between the pattern of neural activity and the different stimuli; 1 split (testing set) was used to make predictions about which stimulus was delivered given the pattern of activity. To compute the classification accuracy, this process was repeated 10 times using different testing and training sets each time. The classification accuracy is defined as the fraction of trials during each bin that the classifier correctly predicted the stimulus.

### Mixture-selectivity index

A mixture-selectivity index (MSI) was used to quantify the difference in firing rate (−2 to 5 s; 200 ms bins) between a neuron’s response to an odor-taste mixture (e.g., isoamyl acetate-sucrose) and its response to the odor component alone (e.g., isoamyl acetate) and its response to the taste component alone (e.g., sucrose). This analysis tested 8 mixture-stimulus differences (4 Mixture-Odor and 4 Mixture-Taste) for each chemoselective neuron (n=85) for a total of 680 mixture-stimulus responses. A response was considered mixture-selective when the evoked MSI score exceeded the mean baseline MSI + 6 x standard deviation. The absolute difference in MSI was used to calculate the average MSI time course (Fig. 5B) to account for the differences between mixtures and components irrespective of excitation or suppression. Significant changes from baseline in the average MSI time course were determined using a Wilcoxon rank sum test with correction for family-wise error (two consecutive significant bins, *P* < 0.05).

### Palatability index

The palatability index (PI) was used to evaluate whether the activity of neurons in the mediodorsal thalamus represents palatability-related features of tastes (Fontanini et al., 2009; Piette et al., 2012; Jezzini et al., 2013; Liu and Fontanini, 2015; Samuelsen and Fontanini, 2017; Bouaichi and Vincis, 2020). This analysis quantifies differences in activity between tastes with similar hedonic values (sucrose/NaCl, citric acid/quinine) and tastes with opposite hedonic values (sucrose/quinine, sucrose/citric acid, NaCl/quinine, NaCl/citric acid). To control for differences in firing rates, the auROC normalized activity (−2 to 5 s, 200 ms bins) was used to estimate the differences between taste pairs. The PI is defined as the difference in the absolute value of the log-likelihood ratio of the auROC normalized firing rate for taste responses with opposite (<|LR|>_opposite_) and similar (<|LR|>_same_) hedonic values. The PI is defined as follows (<|LR|>_opposite_ − <|LR|>_same_), where:

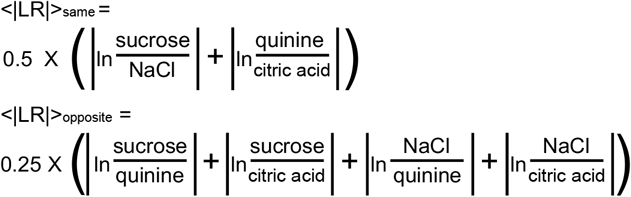

A positive PI value indicates that a neuron responds similarly to tastes with similar palatability and differently to stimuli with opposite hedonic values. A chemoselective neuron was deemed palatability-related when the evoked PI value was positive and exceeded the mean + 6 X standard deviation of the baseline. Significant changes from baseline in the average PI time course were determined using a Wilcoxon rank sum test with correction for family-wise error (two consecutive significant bins, *P* < 0.05).

### Histology

After recordings were completed, rats were anesthetized with ketamine/xylazine/ acepromazine mixture (KXA; 100, 5.2, and 1 mg/kg) and DC current (7 µA for 7 s) was applied to mark the tetrode locations. Rats were then transcardially perfused with cold phosphate buffer solution followed by 4% paraformaldehyde (PFA). Brains were extracted, post-fixed in 4% PFA, then incubated in 30% sucrose. Sections were cut 70 µm thick using a cryostat, mounted, and stained with cresyl violet. Tetrode placement within the mediodorsal thalamus was required for recording sessions to be included in the data analysis (see Fig. 1).

**Figure 1.**
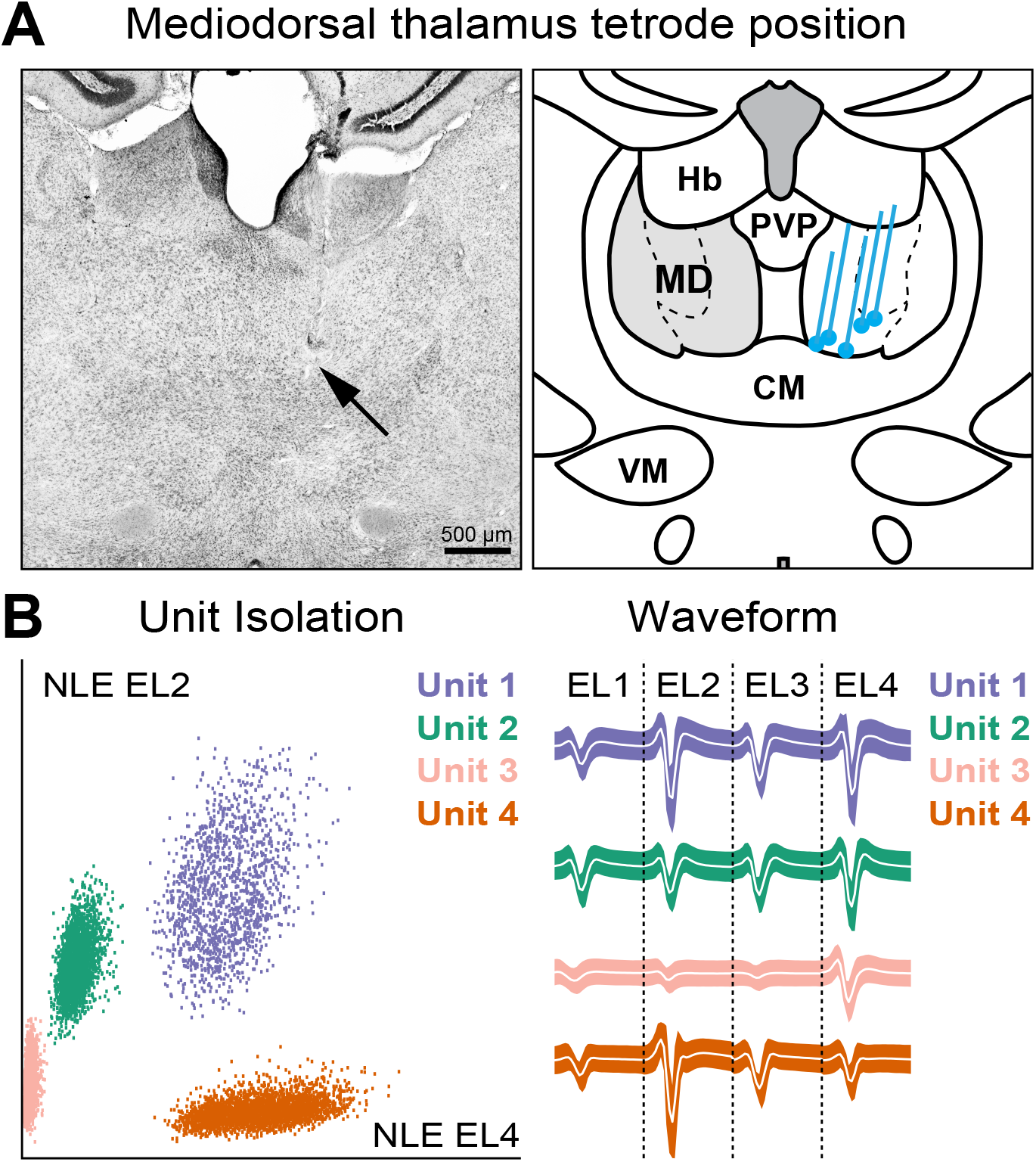
Tetrode locations and representative single-unit recording. ***A***, *Left:* Example histological section showing the recording tetrode position (black arrow) in the mediodorsal thalamus. *Right:* Schematic summary of the reconstructed path of the tetrodes from 5 rats. The blue lines correspond to the dorsoventral range of each drivable tetrode bundle. CM, central medial thalamic nucleus. Hb, habenular nucleus. MD, mediodorsal thalamus. PVP, paraventricular thalamic nucleus. VM, ventromedial thalamic nucleus. ***B***, *Left:* Representative single-unit recordings in the mediodorsal thalamus showing the principal component analysis of waveform shapes of 4 individual neurons. EL, electrode; NLE, Non-Linear energy. *Right:* Average single-unit response for the same 4 neurons recorded from each of the tetrode’s 4 wires.

### Experimental design and statistical analysis

As with previous head-fixed recording experiments (Jones et al., 2007; Fontanini et al., 2009; Samuelsen et al., 2012, 2013; Gardner and Fontanini, 2014; Vincis and Fontanini, 2016; Samuelsen and Fontanini, 2017), only adult female rats were used because the size and strength of adult male rats significantly increases the risk of catastrophic head-cap failure. All chemosensory stimuli were delivered pseudo-randomly by custom written MATLAB (MathWorks) scripts. Experimenters had no control over the order of stimulus delivery. All statistical analyses were performed with GraphPad Prism (GraphPad Software, San Diego, CA) and MATLAB (MathWorks), including population decoding analyses using the neural decoding toolbox (Meyers, 2013). No statistical methods were used to predetermine sample sizes, but the number of recorded neurons and animals in this study are similar to those reported in the field. Significant change from baseline was determined using a Wilcoxon rank-sum comparison between baseline bin and evoked bins with correction for family-wise error (two consecutive significant baseline, *P* < 0.05). Significant differences evoked by the twelve intraoral stimuli were determined using a two-way ANOVA with a Sidak correction for family-wise error ([stimulus X time], main effect of stimulus, *P* < 0.01). Proportional analyses were performed using a *X*^2^ (*P* < 0.05). Post hoc comparisons were made using Fisher’s exact test with Dunn–Sidak correction for familywise error. Kolmogorov–Smirnov (K–S) tests were used to compare distributions of continuous data. Comparisons of response latency and duration between stimulus categories were made using the Kruskal-Wallis test with Tukey HSD correction for family-wise error.

## Results

Previous electrophysiological studies show that neurons in the mediodorsal thalamus represent the identity of odors sampled via orthonasal olfaction (Yarita et al., 1980; Imamura et al., 1984; Courtiol and Wilson, 2014, 2016), but it is unclear how chemosensory signals originating from the mouth are processed by the mediodorsal thalamus. To determine how neurons in the mediodorsal thalamus represent different categories of chemosensory stimuli, we recorded single-unit activity during the intraoral delivery of distilled water, individual odors, individual tastes, and odor-taste mixtures. Figure 1 shows a representative example and a schematic illustration of the dorsal-ventral range of each animal’s recording electrodes in the mediodorsal thalamus. A total of 135 single neurons were recorded from 5 rats across 27 sessions (5.4 ± 0.4 sessions per rat) with an average yield of 5.1 ± 0.9 neurons per session.

### Neurons in the mediodorsal thalamus dynamically represent intraoral chemosensory signals

As a first step to evaluating the neural dynamics evoked by different categories of intraoral chemosensory stimuli, we identified the population of neurons in the mediodorsal thalamus that respond differently to odors, tastes, and odor-taste mixtures (i.e., chemoselective). For a neuron to be defined as ‘chemoselective’, it had to exhibit a significant change from baseline and respond differently across the intraoral stimuli (see Materials and Methods for details). This double criterion was purposefully stringent because the intraoral delivery of solutions could introduce potential confounds related to general effects of somatosensation or attention rather than chemosensory-related activity. We found that 63% (85/135) of the neurons recorded from the mediodorsal thalamus met both criteria and focused our analyses on this chemoselective population. Figure 2A shows the chemoselective population’s average normalized response (population PSTHs) to water and the three odors (top), the four tastes (middle), and the four odor-taste mixtures (bottom). The volatility of the population PSTHs suggested response heterogeneity between stimuli and across time. These differences in response dynamics are illustrated by two representative chemoselective neurons in Figure 2B, where activity was excited (Fig. 2B, left) or suppressed (Fig. 2B, right) by the intraoral delivery of chemosensory stimuli.

**Figure 2.**
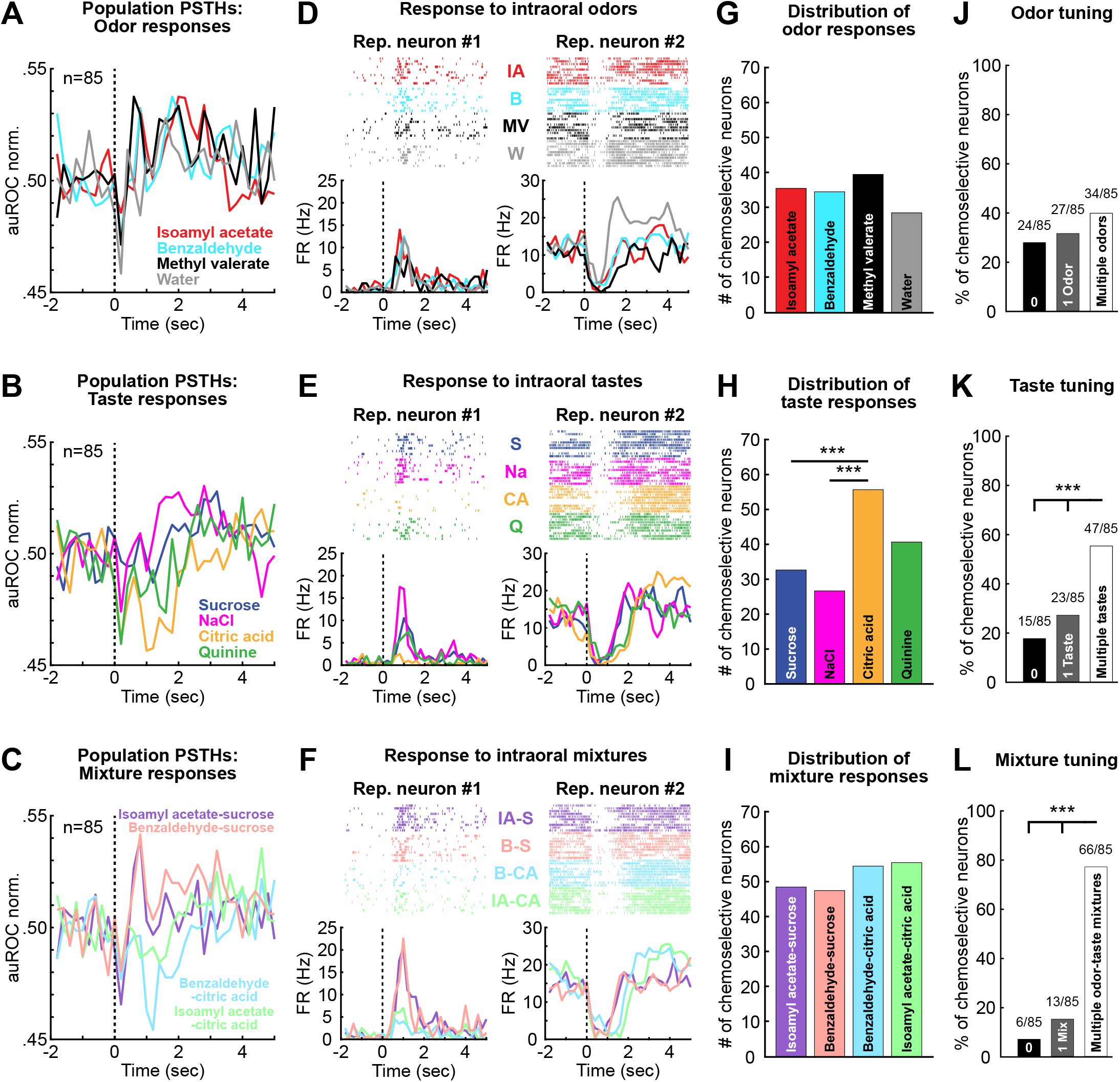
Neurons in the mediodorsal thalamus represent chemosensory signals originating in the mouth. ***A-C***, Peristimulus time histograms (PSTHs) of the chemoselective population’s (n=85) normalized response (auROC; area under the receiver-operating characteristic) to **A**. odors, **B**. tastes, and **C**. odor-taste mixtures. ***D-F***, representative chemoselective neurons firing rate raster plots and PSTHs to the intraoral delivery (time = 0, vertical dashed line) of **D**. water and the three odors (isoamyl acetate [red], benzaldehyde [cyan], methyl valerate [black], water [gray]), **E**. the four tastes (sucrose [blue], NaCl [magenta], citric acid [yellow], quinine [green]), and **F**. the four odor-taste mixtures (isoamyl acetate-sucrose [purple], benzaldehyde-sucrose [peach], benzaldehyde-citric acid [light blue], isoamyl acetate-citric acid [light green]). The representative examples illustrate the heterogeneity in responses, where activity was excited (left) or suppressed (right) by the intraoral delivery of chemosensory stimuli. ***G-I***, Distribution of the number of chemoselective neurons responding to **G**. water and the three odors, **H**. the four tastes, and **I**. the four odor-taste mixtures. ***J-L***, Tuning profiles within each stimulus category of the chemoselective neurons shows the proportion of neurons that did not respond, responded to a single stimulus, or responded to multiple stimuli for **J**. odors, **K**. tastes (middle), and **L**. odor-taste mixtures. *** *P* < 0.001.

One possible factor contributing to the volatility in population responses is differences in the number of neurons that respond to each chemosensory stimulus. We found that most chemoselective neurons respond to stimuli from all three categories (60.0%, 51/85) with significantly smaller proportions responding to a single stimulus category (Fisher’s exact test, *P* < 0.0001; tastes: 3.5%, 3/85; odors: 2.4%, 2/85; odor-taste mixtures: 7.1%, 6/85) or to two stimulus categories (Fisher’s exact test, *P* < 0.0001; tastes and odor-taste mixtures: 17.6%, 15/85; odors and odor-taste mixtures: 8.2%, 7/85; tastes and odors: 1.2%, 1/85). Next, we examined the proportion of chemoselective neurons that responded to each chemosensory stimulus. Figure 2C shows the distribution of neurons that responded to water and each of the three odors (Fig. 2C, top), the four tastes (Fig. 2C, middle), and the four odor-taste mixtures (Fig. 2C, bottom). Overall, there was a significant difference in the proportion of neurons responding to the various stimuli (*X*^2^ (11) = 56.99, *P* < 0.0001). To determine whether chemosensory stimuli activated different proportions of chemoselective neurons, we compared the proportion responding to water (a non-chemosensory stimulus) to the proportion responding to each chemosensory stimulus. Compared to water (34.2%, 29/85), significantly greater proportions of chemoselective neurons responded to citric acid (65.9%, 56/85; Fisher’s exact test, *P* < 0.001) and each of the odor-taste mixtures: isoamyl acetate-sucrose (57.6%, 49/85; Fisher’s exact test, *P* = 0.01), benzaldehyde-citric acid (64.7%, 55/85; Fisher’s exact test, *P* < 0.001), isoamyl acetate-citric acid (65.9%, 56/85; Fisher’s exact test, *P* < 0.001), and benzaldehyde-sucrose (56.5%, 48/85; Fisher’s exact test, *P* < 0.01). Analysis of the distribution of neurons that responded to chemosensory stimuli within each category (Fig. 2C) showed no difference in the proportion of neurons responding within the odor category (*X*^2^ (2) = 0.028, *P* = 0.986) or odor-taste mixture category (*X*^2^ (3) = 2.477, *P* = 0.480), but showed a significant difference across the taste stimuli (*X*^2^ (3) = 22.38, *P* < 0.001). Post-hoc analyses showed that significantly more neurons responded to citric acid (65.9%, 56/85) than sucrose (38.8%, 33/85; Fisher’s exact test, *P* < 0.001) or salt (31.7%, 27/85; Fisher’s exact test, *P* < 0.001), but not quinine (48.2%, 41/85; Fisher’s exact test, *P* > 0.05).

Next, we determined the tuning properties of the chemoselective neurons and found that the greatest proportion responded to at least one odor-taste mixture (92.9%, 79/85), followed by at least one taste (82.4%, 70/85), and then at least one odor (71.8%, 61/85). We then evaluated the tuning profiles of chemoselective neurons within each stimulus category to determine the proportion of neurons that responded to only a single stimulus (i.e., narrowly-tuned) or multiple stimuli (i.e., broadly-tuned). We found that a significantly higher proportion of chemoselective neurons responded to multiple tastes and odor-taste mixtures, but not multiple odor stimuli (Figure 2D). There was no difference in the proportion of chemoselective neurons that did not respond to odors (28.2%, 24/85), responded to a single odor (31.8%, 27/85), or responded to multiple odors (40.0%, 34/85) (*X*^2^ (2) = 2.788, *P* = 0.25) (Fig. 2D, top). Also, there was no difference between the proportion of neurons that responded to only a single odor (31.8%, 27/85), just two odors (21.2%, 18/85), or to all three odors (18.8%, 16/85; *X*^2^ (2) = 4.439, *P* = 0.11). A greater proportion of chemoselective neurons responded to multiple taste stimuli (55.3%, 47/85) compared to those that responded to only a single taste (27.1%, 23/85; Fisher’s exact test, *P* < 0.001) or did not respond to tastes (17.6%,15/85; Fisher’s exact test, *P* < 0.001) (Fig. 2D, middle). However, there was no difference between the proportion of neurons responding to just one taste (27.1%, 23/85), two tastes (24.7%, 21/85), three tastes (14.1%, 12/85), or all four tastes (16.5%, 14/85; *X*^2^ (3) = 6.116, *P* = 0.11). A greater proportion of chemoselective neurons responded to multiple odor-taste mixtures (77.6%, 66/85) compared to those that responded to only a single odor-taste mixture (15.3%, 13/85; Fisher’s exact test, *P* < 0.001) or did not respond to mixtures (7.1%, 6/85; Fisher’s exact test, *P* < 0.001) (Fig. 2D, bottom). There was no difference between the proportion of neurons responding to just one odor-taste mixture (15.3%, 13/85), two mixtures (30.6%, 26/85), three mixtures (20.0%, 17/85), or all four mixtures (27.1%, 23/85; *X*^2^ (3) = 5.128, *P* = 0.16). Together, these analyses revealed that most neurons in the mediodorsal thalamus are broadly-tuned and selectively represent unimodal and multimodal chemosensory signals, indicating that individual neurons process sensory information across a range of chemosensory stimuli.

### Temporal processing of chemosensory signals by the mediodorsal thalamus

Chemosensory processing in the olfactory and gustatory system is characterized by dynamic and time-varying modulations in activity (Katz et al., 2001; Fontanini et al., 2009; Maier et al., 2012; Samuelsen et al., 2012, 2013; Liu and Fontanini, 2015; Maier, 2017; Samuelsen and Fontanini, 2017). While most of these areas primarily respond with excitation to chemosensory stimuli, a study by Liu and Fontanini (2015) examining another thalamic nucleus, the gustatory thalamus (i.e., the parvicellular portion of the ventroposteromedial nucleus, VPMpc), revealed a near balance between taste-evoked excitation and suppression. Therefore, our first step in examining the neural dynamics of chemosensory-evoked activity in the mediodorsal thalamus was to sort responses into excited responses (when the significant evoked activity was greater than baseline), suppressed responses (when the significant evoked activity was less than the baseline), and non-responsive (those that did not significantly differ from baseline). The population averages of excited and suppressed responses (Fig. 3A) and the heat maps of each significant response (Fig. 3B) to odors (top), tastes (middle), and odor-taste mixtures (bottom) illustrate the heterogeneity of responses across the chemoselective population. Although chemosensory stimuli more often evoked excitation (27.2%, 254/935) than suppression (23.7%, 222/935), there was no significant difference in the overall proportion of responses excited or suppressed by the different chemosensory stimuli (Fisher’s exact test, *P* = 0.0998). This equivalence in stimulus-evoked excitation and suppression was represented within each stimulus category. There was no difference in the proportion of odor responses that evoked excitation (23.5%, 60/255) or suppression (20.0%, 51/255; Fisher’s exact test, *P* = 0.391), in taste responses that evoked excitation (24.1%, 82/340) or suppression (22.1%, 75/340; Fisher’s exact test, *P* = 0.585), or odor-taste mixture responses that evoked excitation (32.9%, 112/340) or suppression (28.2%, 96/340; Fisher’s exact test, *P* = 0.212).

**Figure 3.**
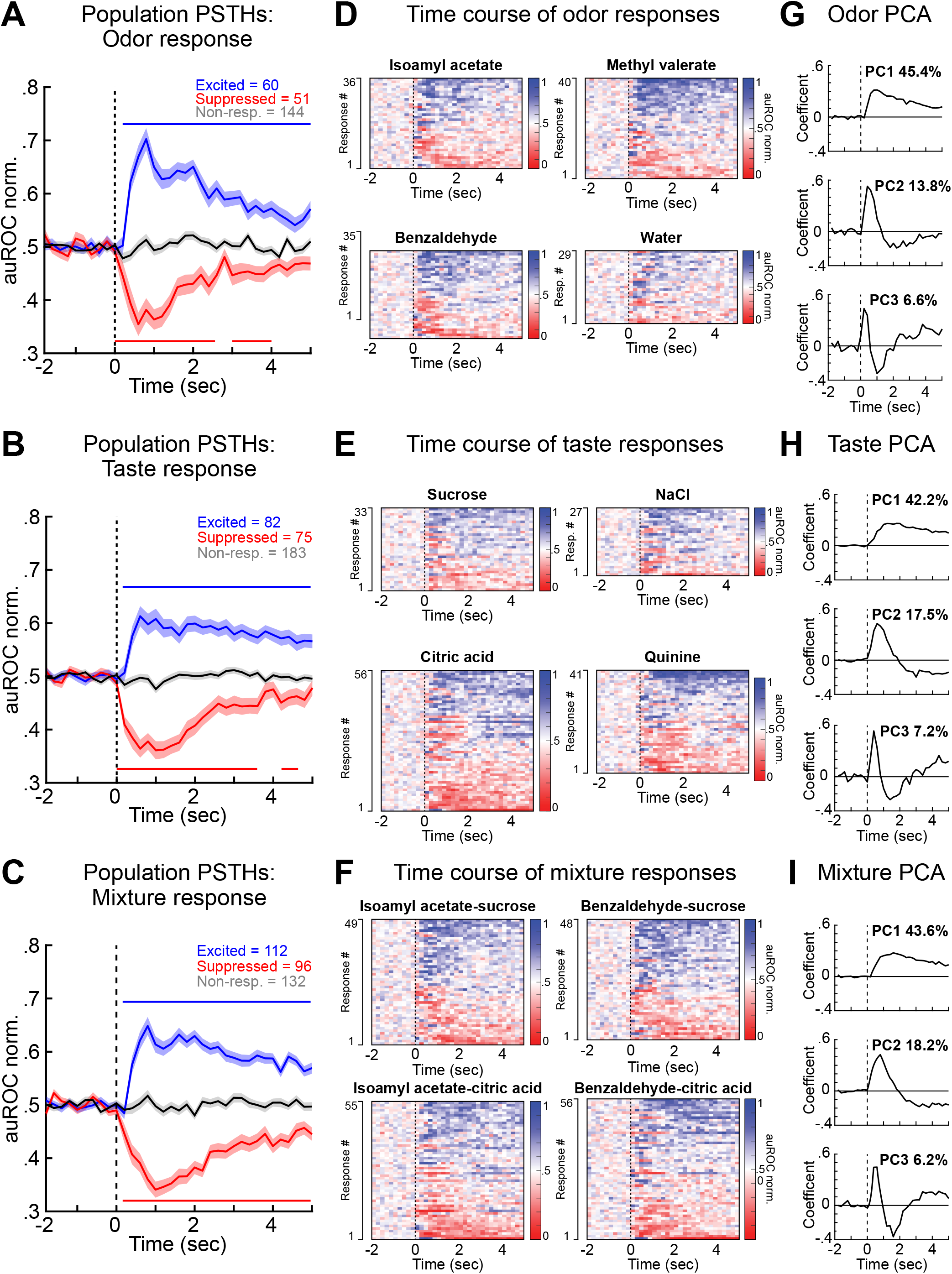
Intraoral chemosensory stimuli evoke excitation and suppression. ***A-C***, auROC normalized population PSTHs of the responses excited (blue), suppressed (red), or non-responses (black) to **A**. odors, **B**. tastes, and **C**. odor-taste mixtures. Vertical dashed line indicates stimulus delivery (time = 0). Shaded area represents the SEM. Horizontal lines above and below traces indicate when responses significantly differed from those that did not respond (Wilcoxon rank sum, p<0.05). ***D-F***, Pseudocolored heat maps of each significant response to **D**. odors, **E**. tastes, and **F**. odor-taste mixtures plotted from the most suppressed to the most excited for each stimulus. ***G-I***, Eigenvectors of the first three principal components of the significant responses to **G**. odors, **H**. tastes, and **I**. odor-taste mixtures. Regardless of the stimulus category, PC 1 represented a monotonic component that lasted the entire 5 s temporal window, PC 2 represented a biphasic component, and PC 3 represented triphasic modulations.

Analysis of the distribution in response latency revealed differences in the onset of excited and suppressed activity. Overall, the onset of responses suppressed by chemosensory stimuli (301.7 ± 30.1 ms) occurred significantly faster than the onset of excitation (443.9 ± 45.0 ms; two-sample Kolmogorov– Smirnov test, K-S stat = 0.23, *P* < 0.001). While there was no difference between the onset of excited responses between the stimulus categories (odors: 443.4 ± 82.8 ms, tastes: 471.6 ± 90.3 ms, odor-taste mixtures: 426.0 ± 65.5 ms; Kruskal-Wallis, H (2) = 0.1080, *P* = 0.948), this analysis revealed a significant difference between stimulus categories for the onset of suppression (Kruskal-Wallis, H (2) = 8.9704, *P* = 0.011). Tukey’s HSD Test for multiple comparisons showed that onset of suppression occurred significantly faster with the intraoral delivery of odor stimuli (156.1 ± 36.1 ms) compared to either tastes (341.0 ± 55.8 ms, *P* = 0.033) or odor-taste mixtures (349.2 ± 49.5 ms, *P* = 0.012).

To better understand the temporal profiles of the chemosensory-evoked activity, we performed a PCA on the auROC normalized activity (−2 to 5s; 200 ms bins) of the significant responses in each stimulus category to extract the most frequent trends in the time course of responses (Narayanan and Laubach, 2009; Liu and Fontanini, 2015) (Fig. 3C). The first three principal components (PCs) accounted for nearly the same total variance of responses to odors (65.8%), tastes (66.9%), and odor-taste mixtures (68%). Furthermore, the responses represented by the three largest eigenvectors were the same across the stimulus categories; where PC 1 represents a monotonic component that lasted the entire 5 s temporal window, PC 2 represents a biphasic component, and PC 3 represents triphasic modulations.

Next, we determined the duration of activity that was significantly greater than baseline (a measure of total excitation) or significantly lower than baseline (a measure of total suppression) differed by stimulus category. This analysis identifies each significant bin over time to account for the heterogeneity of biphasic and triphasic evoked responses (Fig. 3C). Overall, the excited activity lasted significantly longer (716.2 ± 34.7 ms) compared to responses suppressed by chemosensory stimuli (477.2 ± 27.5 ms; two-sample Kolmogorov–Smirnov test, K-S stat = 0.21, *P* < 0.001). However, there were no differences between stimulus categories in the duration of either excited activity (odors: 737.1 ± 74.5 ms, tastes: 742.6 ± 60.3 ms, mixtures: 685.0 ± 51.7 ms; Kruskal-Wallis, H (2) = 0.3723, *P* = 0.8301) or suppressed activity (odors: 528.2 ± 59.2 ms, tastes: 488.6 ± 49.9 ms, mixtures: 441.6 ± 39.3 ms; Kruskal-Wallis, H (2) = 1.4805, *P* = 0.4770). In summary, neurons were suppressed by stimuli more quickly, especially in response to odors, but responded with excitation significantly longer than they were suppressed.

While the activity of individual neurons can represent specific features of chemosensory stimuli, networks of neurons are responsible for integrating and processing that information to guide behavior. We hypothesized that the heterogeneity displayed by the population of chemoselective neurons enables the accurate representation of the various chemosensory stimuli across time. We used a population decoding analysis (Jezzini et al., 2013; Liu and Fontanini, 2015; Bouaichi and Vincis, 2020) to quantify whether the firing patterns of ensembles of chemoselective neurons in the mediodorsal thalamus accurately encodes stimulus identity over time. We computed the decoding performance of the population of chemoselective neurons (n=85) and non-selective neurons (n=50) for the three categories of chemosensory stimuli (Fig. 4). Water was included in the population decoding analysis for each of the three chemosensory categories as a general non-chemosensory stimulus. Figure 4A shows the time course of the classification accuracy for odors and water. Odor decoding of the chemoselective population activity showed an early onset (classification above chance from the first bin after intraoral delivery) before peaking 1 s after odor delivery. The classification accuracy briefly returns to chance level ~3.75 s after stimulus delivery and then moves above chance for the remaining temporal window. The odor classification accuracy of the non-selective neurons only exceeded chance for a single bin at 2 s. Figure 4B shows the time course of the classification accuracy for tastes and water. Taste decoding of the chemoselective population activity did not perform above chance until the second bin (~500 ms), peaked at 1.5 s after intraoral delivery, and remained above chance for the entire 5 s time frame. The taste classification accuracy of the non-selective neurons never exceeded chance. Figure 4C shows the time course of classification accuracy for odor-taste mixtures and water. Like the decoding performance of odors, the odor-taste mixture decoding of the chemoselective population activity showed an early onset (first bin, ~250 ms). The odor-taste mixture classification accuracy peaked 1.25 s after intraoral delivery, slightly after the peak for odors, but before the peak for tastes. Like the decoding performance of tastes, the classification accuracy stayed above chance for the entire 5 s time frame. The odor-taste mixture classification accuracy of the non-selective neurons did not exceed chance but for a single bin at 1.75 s. The decoding performance for odor-taste mixtures is particularly interesting because the four odor-taste mixtures are combinations of just two tastes (sucrose and citric acid) and two odors (isoamyl acetate and benzaldehyde). Together, these data indicate that ensembles of neurons in the mediodorsal thalamus reliably encode unimodal and multimodal chemosensory signals.

**Figure 4.**
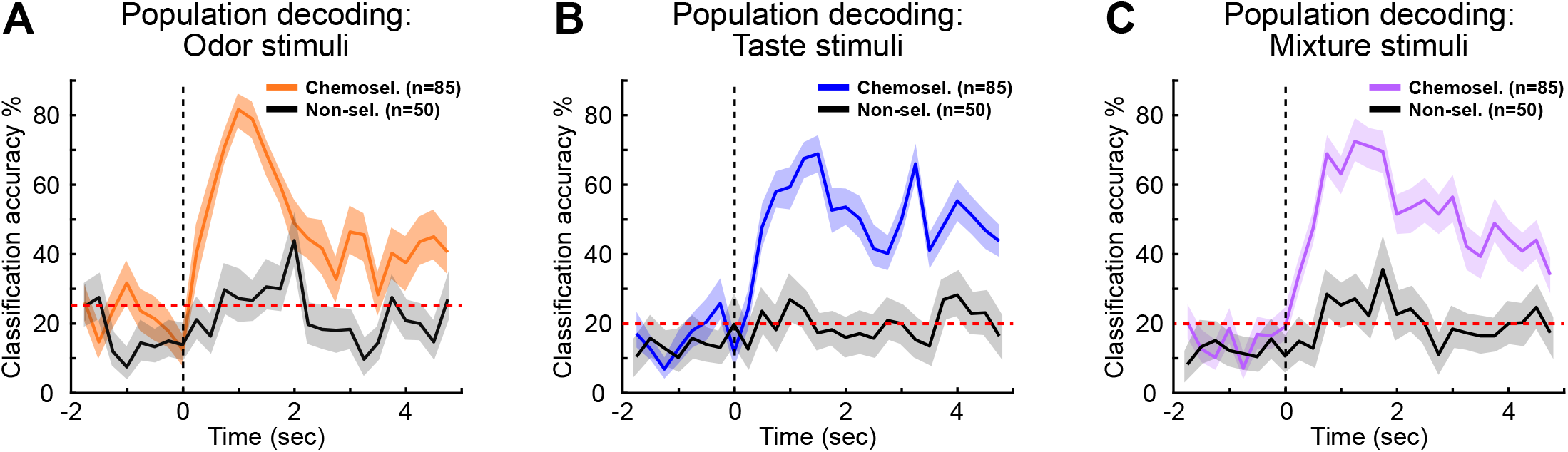
The population decoding performance over time of the chemoselective neurons (n = 85) and non-selective neurons (n = 50) for each category of chemosensory stimuli for ***C***, odors, ***D***, tastes, and ***E***, odor-taste mixtures. Note that the chemoselective population activity represented odors and odor-taste mixtures more quickly than tastes, but sustained responses to tastes and odor-taste mixtures longer than odors. Red-dashed line indicates chance-level. Vertical dashed line indicates stimulus delivery (time = 0). The shaded area represents a 99.5% bootstrapped confidence interval (CI).

### A subset of chemoselective neurons represents mixtures differently from its components

Although most chemoselective neurons respond to odor-taste mixtures, they could be responding to the odor or taste component of the mixture. If so, one would expect that the activity evoked by an odor-taste mixture to be similar to the response elicited by its odor or taste component alone. Visual inspection of the raster plots and PSTHs of individual neurons in the mediodorsal thalamus indicated that some responded differently to odor-taste mixtures compared to their individual odor or taste components (Fig. 5A). We used a mixture-selectivity index (MSI) (see Materials and Methods for details) to examine whether neurons in the mediodorsal thalamus respond to odor-taste mixtures differently than to their unimodal components. This analysis quantifies the difference in firing rate across time between a neuron’s response to an odor-taste mixture (e.g., isoamyl acetate-sucrose) and the response to its odor component alone (e.g., isoamyl acetate) or its taste component alone (e.g., sucrose). A response was considered significantly different when the evoked MSI score exceeded the mean baseline MSI + 6 x standard deviation.

**Figure 5.**
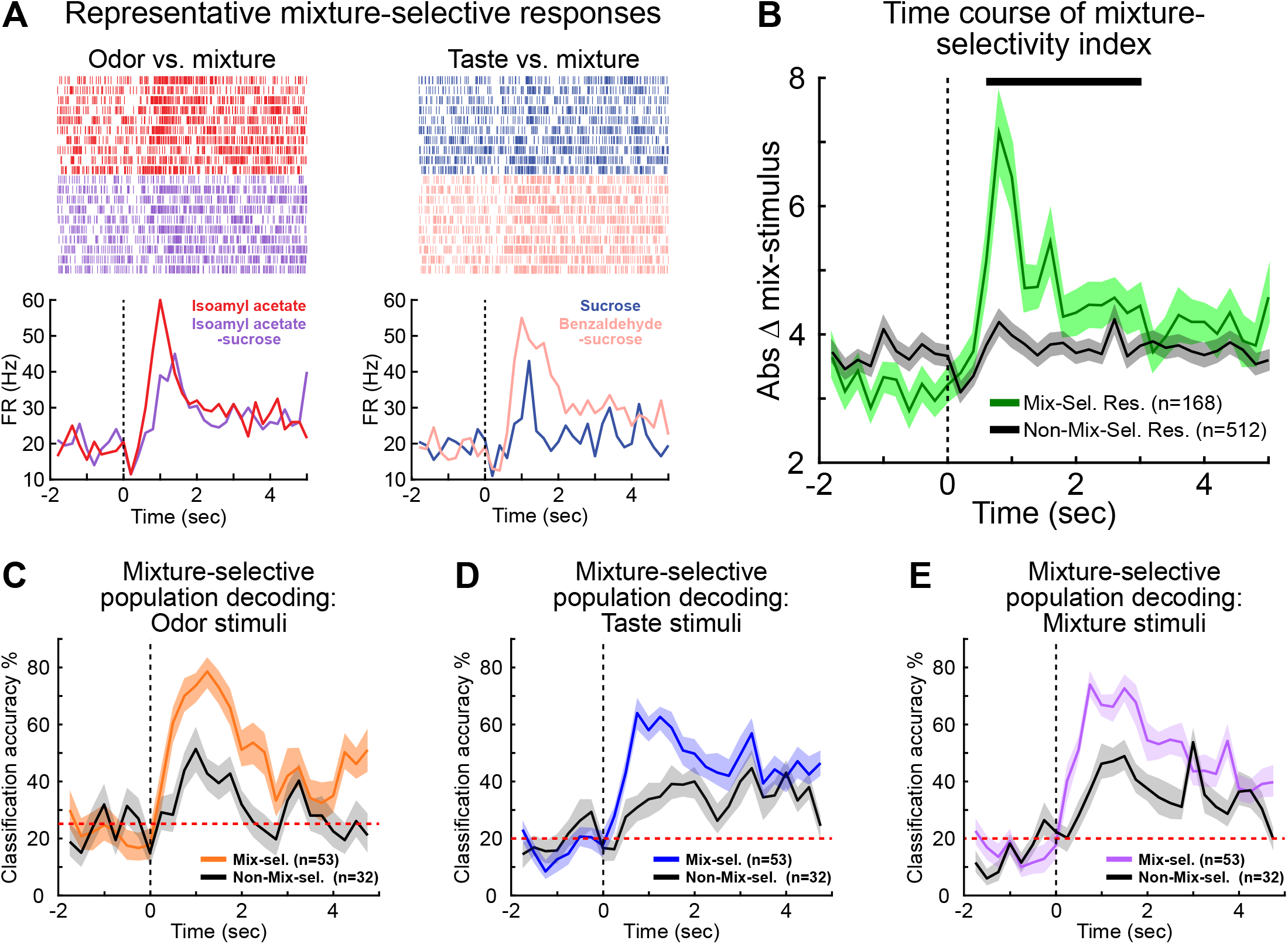
Mixture-selectivity index (MSI) of neurons in the mediodorsal thalamus. ***A***, Raster plots and PSTHs from a representative neuron in the mediodorsal thalamus illustrating the differences in activity evoked by mixtures and their components: isoamyl acetate-sucrose mixture vs. isoamyl acetate alone (left) and benzaldehyde-sucrose mixture and sucrose alone (right). Vertical dashed line indicates stimulus delivery (time = 0). ***B***, Time course of the average absolute mixture-selectivity index values for the 168 mixture-selective responses (green line) and the 512 non-mixture-selective responses (black line) −2 s before to 5 s after intraoral delivery (200 ms bins). The mixture-selective responses significantly differ from baseline from 0.6 – 3 s (black bar) after stimulus delivery, while the non-mixture-selective responses never differ from baseline. Shaded area represents the SEM. ***C-E***, Population decoding performance over time of mixture-selective (n = 53) and non-mixture-selective neurons (n = 32) for ***C***, odors, ***D***, tastes, and ***E***, odor-taste mixtures. Note that both populations had decoding performances above chance (red-dashed line) but with different temporal profiles. Black vertical dashed line indicates stimulus delivery (time = 0). The shaded area represents a 99.5% bootstrapped confidence interval (CI).

The MSI analysis revealed that nearly a third of the odor-taste mixture responses (32.8%, 168/680) differed from at least one of its components. These 168 mixture-selective responses could be represented by as few as 21 neurons or they could be spread across the entire chemoselective population because each neuron could account for a maximum of 8 mixture-selective responses (each of the 4 odor-taste mixtures compared to their odor and taste component). However, we found that 53 of the 85 (62.4%) chemoselective neurons accounted for the mixture-selective responses. Of the 53 neurons, 21 (39.6%, 21/53) had odor-taste mixture responses that differed from odor and taste responses, 23 (43.4%, 23/53) differed from just odor responses, and 9 (17.0%, 9/53) differed from just taste responses. To examine these differences over time, we calculated the average absolute difference in MSI (−2 to 5s; 200 ms bins) for both the mixture-selective and non-mixture-selective responses (Fig. 5B). The absolute difference in MSI was used to account for differences between odor-taste mixtures and components irrespective of excitation or suppression. This analysis revealed that the mixture-selective responses began to significantly differ from baseline 600 ms after stimulus delivery, peaked at 800 ms, and remained significantly above baseline for another 2.2 s (green line, black bar, Wilcoxon rank-sum, two consecutive significant bins, *P* < 0.05). As a control, the absolute value of the MSI was calculated for the non-mixture-selective responses. There was no difference from baseline (black line, *P* > 0.05).

Next, we used the population decoding analysis to quantify the contribution of the ensemble of mixture-selective neurons to the representation of chemosensory stimuli over time. Figure 5 shows the decoding performance of the population of mixture-selective neurons (n=53) and the non-mixture-selective neurons (n=32) for the three categories of chemosensory stimuli. Importantly, both groups had decoding performances above chance but with different temporal profiles. The decoding of the mixture-selective population activity showed an early onset, with classification accuracy above chance from the first bin, for all three chemosensory categories. While classification accuracy of the non-mixture-selective population did not exceed chance until 500ms for tastes (Fig. 5B) and odor-taste mixtures (Fig. 5C), and 750 ms for odors (Fig. 5A). Although the classification accuracy of both groups remained above chance when decoding tastes and odor-taste mixtures, the decoding performance for odors differed between the mixture-selective and non-mixture-selective populations. The classification accuracy of the mixture-selective neurons remained above chance for the entire period, but the classification accuracy of the non-mixture-selective population returned to chance 2 s after odor delivery (Fig. 5A). These results suggest that differences between odor-taste mixtures and their unimodal components are distributed across the population of chemoselective neurons in the mediodorsal thalamus.

### A subset of chemoselective neurons represents taste palatability

Tastes have intrinsic values, with rodents consuming palatable tastes and avoiding unpalatable ones. It is well established that brain regions important for chemosensory processing and feeding-related behaviors represent the chemical and hedonic properties of tastes (Fontanini et al., 2009; Piette et al., 2012; Sadacca et al., 2012; Jezzini et al., 2013; Li et al., 2013; Liu and Fontanini, 2015; Samuelsen and Fontanini, 2017). However, its unknown whether the activity in the mediodorsal thalamus represents taste palatability, meaning that tastes belonging to similar hedonic categories evoke similar responses (e.g., sucrose/NaCl vs. citric acid/quinine). Therefore, we calculated a palatability index (PI) (see Materials and Methods for details) to determine whether neurons in the mediodorsal thalamus represent palatability-related features of tastes. This analysis quantifies the differences in activity between tastes of similar palatability (sucrose/NaCl, citric acid/quinine) and tastes of opposite palatability (sucrose/quinine, sucrose/citric acid, NaCl/quinine, NaCl/citric acid). A chemoselective neuron was considered to represent taste palatability when it had a positive PI value (it responded similarly to tastes with similar hedonic value but differently to tastes with opposite hedonic value) and the evoked PI value exceeded the mean + 6 x standard deviation of the baseline.

This analysis revealed that the activity of more than a quarter (27.1%, 23/85) of the chemoselective neurons represented taste palatability (Fig. 6A, representative examples). The average PI value (−2 to 5s; 200 ms bins) of the palatability-related (n = 23) and non-palatability neuron populations (n = 62) was used to examine the temporal evolution of palatability-related activity. Figure 6B shows that the mean PI value of the palatability-related neurons began to significantly differ from baseline beginning at 1.4 s and peaked 2 s after stimulus delivery (green line, black bars, Wilcoxon rank-sum, two consecutive significant bins, *P* < 0.05). The mean response of the non-palatability population did not differ from baseline (black line, *P* > 0.05).

**Figure 6.**
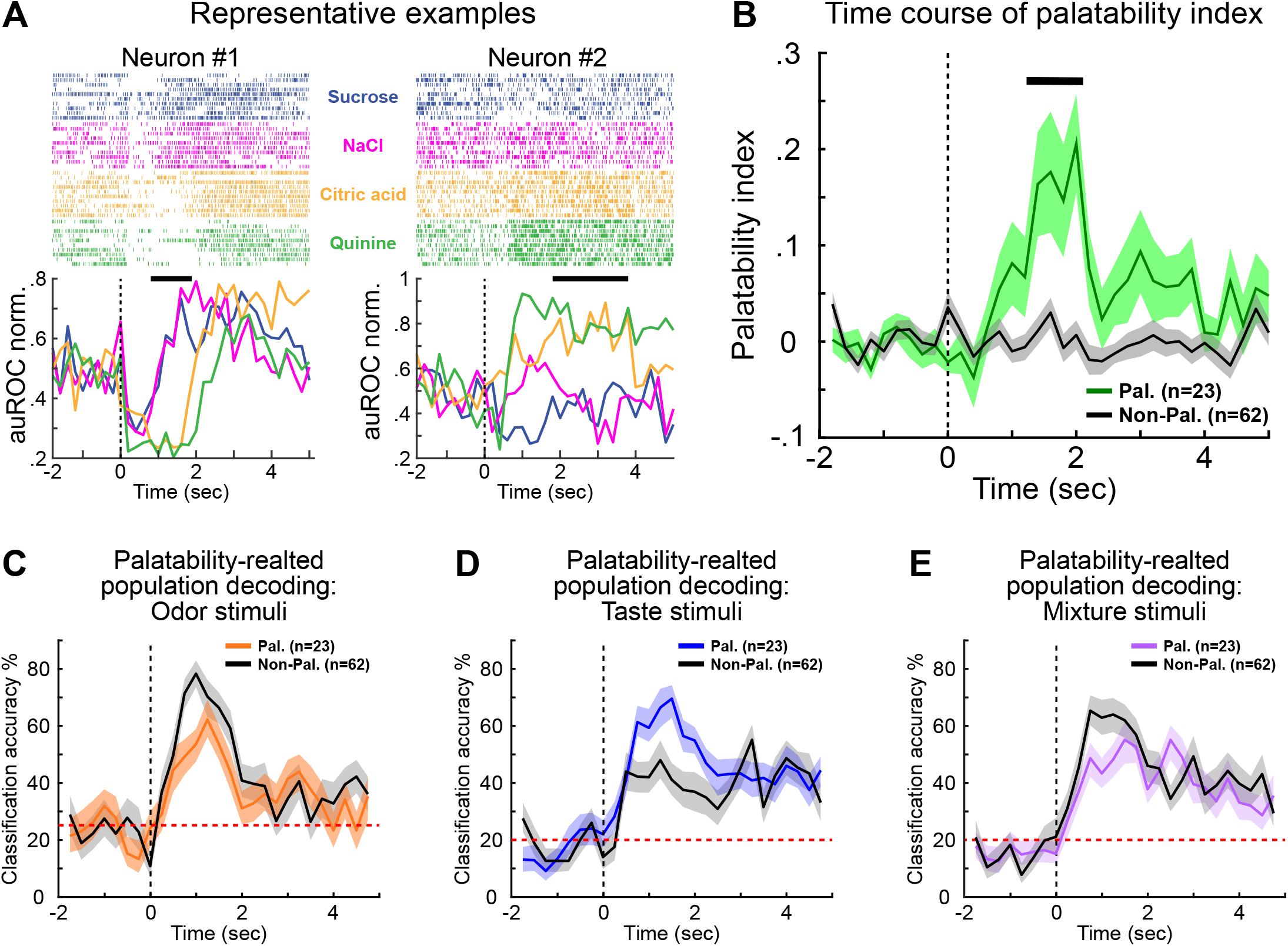
Processing of taste palatability by neurons in the mediodorsal thalamus. ***A***, Raster plots and PSTHs from two neurons in the mediodorsal thalamus that represent the palatability-related features of tastes. The horizontal black bars indicate significant palatability-related difference in activity. Vertical dashed line indicates stimulus delivery (time = 0). ***B***, Time course of the average palatability index (PI) value of the 23 palatability-related neurons (green line) and 62 non-palatability neurons (black line) −2 s before to 5 s after intraoral delivery (200 ms bins). The response of the palatability-related population significantly differs from baseline from 1.4 – 2 s (black bar) after stimulus delivery, while the average PI value of the non-palatability population never differs from baseline. Vertical dashed line indicates stimulus delivery (time = 0). Shaded area represents the SEM. ***C-E***, Population decoding performance over time of palatability-related (n = 23) and non-palatability neurons (n = 62) for ***C***, odors, ***D***, tastes, and ***E***, odor-taste mixtures. Note that both populations performed better than chance (red-dashed line) but with categorical and temporal differences. Vertical dashed line indicates stimulus delivery (time = 0). The shaded area represents a 99.5% bootstrapped confidence interval (CI).

It is possible that this palatability-related population of chemoselective neurons primarily represents taste signals and does not carry information relative to odors or odor-taste mixtures. Therefore, a population decoding analysis was used to examine how the palatability-related population represents the different categories of chemosensory stimuli (Fig. 6C-E). While both populations performed better than chance for all three categories, this analysis revealed categorical and temporal differences between the palatability-related and the non-palatability chemoselective populations. The palatability-related neurons represented taste information better than the non-palatability chemoselective population, but during a specific temporal window (0.5-2.25 s). The opposite occurred with odors and odor-taste mixtures; the non-palatability chemoselective population performed better than palatability-related neurons for a brief period for both the odors (0.5-1 s) and odor-taste mixtures (0.5-1.25 s). These results indicate that a subset of chemoselective neurons in the mediodorsal thalamus encode the hedonic properties of tastes. However, this population is multidimensional and represents odors and odor-taste mixtures as well.

Taken together, our results show that neurons in the mediodorsal thalamus represent the sensory properties of odors, tastes, and odor-taste mixtures with information relative to mixture-selectivity and taste palatability represented by the neural activity. These findings suggest that neurons in the mediodorsal thalamus are capable of representing the sensory and hedonic properties of unimodal and multimodal chemosensory stimuli important for making decisions about food.

## Discussion

Higher-order thalamic areas are thought to modulate, synchronize, and transmit behaviorally relevant information between sensory and higher-order cortical areas (Theyel et al., 2010; Saalmann et al., 2012; Stroh et al., 2013; Mease et al., 2016; Zhou et al., 2016; Schmitt et al., 2017; Rikhye et al., 2018). By sustaining communication across cortical regions, these cortico-thalamo-cortical (i.e., transthalamic) circuits separate potentially overlapping information and enable rapid behavioral changes based on environmental demands (Saalmann, 2014; Sherman, 2016; Rikhye et al., 2018). Given its connectivity, the mediodorsal thalamus may perform a similar function by communicating behaviorally-relevant chemosensory information between principal regions of the olfactory and gustatory systems and higher-order cortical areas (Price and Slotnick, 1983; Kuroda et al., 1992; Ray and Price, 1992; Shi and Cassell, 1998; Kuramoto et al., 2017; Pelzer et al., 2017). The results presented here are the first to demonstrate how neurons in the mediodorsal thalamus represent olfactory and gustatory signals originating from the mouth. Tetrode recordings in behaving rats revealed that most neurons in the mediodorsal thalamus dynamically represent intraoral odors, tastes, and odor-taste mixtures. These chemoselective neurons responded broadly across intraoral stimuli with time-varying multiphasic changes in activity split between excitation and suppression. Population analyses revealed temporal differences in responses to chemosensory stimuli, where chemoselective neurons responded to odors and odor-taste mixtures more quickly than tastes, but sustained responses to tastes and odor-taste mixtures longer than odors. Analyses of the temporal sequence of mixture-selectivity and taste palatability revealed that information related to chemical identity is represented before taste palatability. Our results further demonstrate the multidimensionality of the mediodorsal thalamus and provides additional evidence of its involvement in processing chemosensory information important for consummatory behaviors.

Similar to the findings of electrophysiological studies of the gustatory thalamus (Liu and Fontanini, 2015), basolateral amygdala (Fontanini et al., 2009), piriform cortex (Maier et al., 2012; Maier, 2017), gustatory cortex (Katz et al., 2001; Samuelsen et al., 2012, 2013; Samuelsen and Fontanini, 2017), and medial prefrontal cortex (Jezzini et al., 2013), chemosensory processing by the mediodorsal thalamus is characterized by dynamic and time-varying modulations in activity. We found that 63% of recorded neurons responded selectively to passively delivered intraoral stimuli, with similar proportions of this chemoselective population responding to odors, tastes, and odor-taste mixtures. Analysis of the breadth of tuning revealed that chemoselective neurons responded broadly to tastes and odor-taste mixtures, but 44% of odor-responsive neurons responded to just a single odor. Previous electrophysiological studies report varying degrees of odor specificity in the mediodorsal thalamus. For example, recordings in anesthetized rabbits showed that 24% of neurons responded to a single orthonasal odor (Imamura et al., 1984), while recordings in anesthetized rats found that 63% of neurons responded to a single orthonasal odor (Courtiol and Wilson, 2014). These findings suggest that the degree of odor specificity in the mediodorsal thalamus likely depends on a variety of factors, including the size of the odor set, route of delivery, and state of the animal.

It is well established that neurons in the mediodorsal thalamus respond to visual, auditory, somatosensory, and olfactory stimuli (Yarita et al., 1980; Imamura et al., 1984; Oyoshi et al., 1996; Yang et al., 2006; Courtiol and Wilson, 2016), but, to the best of our knowledge, only one study has provided evidence of taste-evoked activity in the mediodorsal thalamus. Oyoshi and colleagues (1996) showed that neurons respond to sucrose when it was given as a reward for the correct choice in a sensory-discrimination task. However, given the complex nature of the task design, it is unclear whether responses were somatosensory-, reward-, or taste-dependent. The results presented here clearly demonstrate that neurons in the mediodorsal thalamus represent the sensory and hedonic properties of taste stimuli. Additionally, our findings reveal that intraoral olfactory and gustatory signals converge onto individual neurons in the mediodorsal thalamus. Convergence of chemosensory signals is not unique to the mediodorsal thalamus as the piriform cortex and gustatory cortex are known to respond to both odor and taste stimuli (Maier et al., 2012; Maier, 2017; Samuelsen and Fontanini, 2017). However, the convergence of olfactory and gustatory signals in the mediodorsal thalamus may be specific to the chemosensory modalities, since previous studies found that odor-responsive neurons did not respond to other sensory modalities, (e.g., visual, auditory, or somatosensory stimuli) (Yarita et al., 1980; Imamura et al., 1984).

To challenge multimodal chemosensory processing by the mediodorsal thalamus, we chose to use a four odor-taste mixture set comprised of just two tastes (sucrose and citric acid) and two odors (isoamyl acetate and benzaldehyde). If neurons primarily represented either unimodal odor or taste signals, the neural decoder would be unable to distinguish between mixtures containing the same odor or the same taste. The population decoding performance showed that chemoselective neurons accurately classified odors, tastes, and odor-taste mixtures, suggesting that they differently encode unimodal and multimodal chemosensory stimuli. This was confirmed using a mixture-selectivity analysis, which showed that most chemoselective neurons respond differently to odor-taste mixtures and their individual components beginning ~600ms after stimulus delivery. Taken together, our findings demonstrate that neurons in the mediodorsal thalamus dynamically represent unimodal and multimodal gustatory and olfactory stimuli originating from the mouth.

While identifying the sources of chemosensory information to the mediodorsal thalamus is outside the scope of this study, analysis of the temporal dynamics of thalamic activity indicates likely sources of input. The thalamic representation of chemosensory information is rapid and persistent, but with time-varying differences between categories of stimuli. Our results show that the mediodorsal thalamus encodes the chemical identity of stimuli containing odors (odors and odor-taste mixtures) ~250 ms after intraoral delivery but takes an additional ~250 ms to encode the identity of tastes alone. However, both the piriform cortex and gustatory cortex encode chemical identity well before the mediodorsal thalamus. For example, neurons in the piriform cortex accurately represent odor identity ~100 ms after inhalation (Bolding and Franks, 2017), while neurons in the gustatory cortex encode taste identity ~175-250 ms (Katz et al., 2001; Jezzini et al., 2013; Bouaichi and Vincis, 2020). On the other hand, higher-order cortical areas like the medial prefrontal cortex do not encode taste identity until ~575 ms after intraoral delivery (Jezzini et al., 2013), much later than the mediodorsal thalamus. These findings suggest a mechanism whereby the mediodorsal thalamus initially receives chemosensory information from the piriform cortex and gustatory cortex, while dynamic multiphasic activity arises via recurrent interactions with the chemosensory cortices and higher-order cortical areas. This transthalamic circuit could act as a means for large-scale integration of chemosensory information across multiple cortical circuits (Saalmann, 2014).

Similar network interactions may be responsible for the hedonic representation of tastes by the mediodorsal thalamus. Based on connectivity, the basolateral amygdala and gustatory cortex are the most likely sources of hedonic information to the mediodorsal thalamus. Both areas are densely reciprocally connected with the mediodorsal thalamus and represent taste palatability before the mediodorsal thalamus. The basolateral amygdala encodes taste palatability between ~0.25–1.0 s after taste delivery (Fontanini et al., 2009), while the gustatory cortex doesn’t begin to represent taste palatability until ~0.75-1.0 s (Katz et al., 2001; Jezzini et al., 2013; Samuelsen and Fontanini, 2017). The palatability index time course (Fig. 6) suggests multiple periods of hedonic processing in the mediodorsal thalamus that overlap ~1.5 s after stimulus delivery. The temporal variations in hedonic processing by the mediodorsal thalamus may arise from contributions by the basolateral amygdala to the initial ramp and the gustatory cortex modulating and sustaining palatability-related information. These hypotheses require future studies selectively targeting neuronal populations with cell-specific viral manipulations (e.g., optogenetics) to elucidate the contribution of these regions to chemosensory processing by the mediodorsal thalamus.

In summary, despite its robust connectivity with olfactory and gustatory areas, involvement in olfactory-dependent behaviors, and importance for the perception of flavors, the mediodorsal thalamus remains an understudied area of the network that processes chemosensory information. Our results show that neurons in the mediodorsal thalamus dynamically encode the sensory and hedonic properties of chemosensory signals originating from the mouth. Future studies probing cortico-thalamo-cortical interactions in behaving animals are necessary to determine the contribution of the mediodorsal thalamus for chemosensory-dependent behaviors.

## Acknowledgements

This work was supported by a National Institute of Deafness and Other Communication Disorders Grant (R01-DC018273; CS). The authors would like to thank Dr. Sanaya Stocke for the helpful comments and discussions.

